# The Moose of Isle Royale: An Unnatural Condition?

**DOI:** 10.1101/019984

**Authors:** Samuel V. Scarpino, Rafael F. Guerrero, Philip V. Scarpino

## Abstract

The now iconic moose of Isle Royale National Park arrived on the island sometime between 1910 and 1915 (Hickie, 1936; Murie, 1934). Prior to that period there is no evidence of moose in either naturalist reports or in the archaeological history of the island (Murie, 1934; Scarpino, 2011). Early naturalists–while observing the moose during their first 20 years on the island—noted both their dramatic expansion, and equally dramatic population crash in the 1930s, see Figure 1. Around 1950, and just as the moose were rebounding, wolves crossed a frozen Lake Superior and began what is now one of our most emblematic predator/prey systems (Peterson, 1995). Recently, the wolves on Isle Royale appear headed for local extinction (Mlot, 2015). Calls to repopulate the island have renewed the vigorous debate surrounding what is and what is not wild about Isle Royale (Scarpino, 2011; Cronon, 1996; Nelson & Callicott, 2008; Cronon, 2003; Peterson, 1999).

---

> *Dead and dying moose were frequently found lying in the heavy snow along the lake shores from January until Spring.*
>
> — -Paul F. Hickie, Isle Royale Moose Studies 1936

The now iconic moose of Isle Royale National Park arrived on the island sometime between 1910 and 1915 (Hickie, 1936; Murie, 1934). Prior to that period there is no evidence of moose in either naturalist reports or in the archaeological history of the island (Murie, 1934; Scarpino, 2011). Early naturalists-–while observing the moose during their first 20 years on the island—noted both their dramatic expansion, and equally dramatic population crash in the 1930s, see Figure 1. Around 1950, and just as the moose were rebounding, wolves crossed a frozen Lake Superior and began what is now one of our most emblematic predator/prey systems (Peterson, 1995). Recently, the wolves on Isle Royale appear headed for local extinction (Mlot, 2015). Calls to repopulate the island have renewed the vigorous debate surrounding what is and what is not wild about Isle Royale (Scarpino, 2011; Cronon, 1996; Nelson & Callicott, 2008; Cronon, 2003; Peterson, 1999).

**Figure 1.**
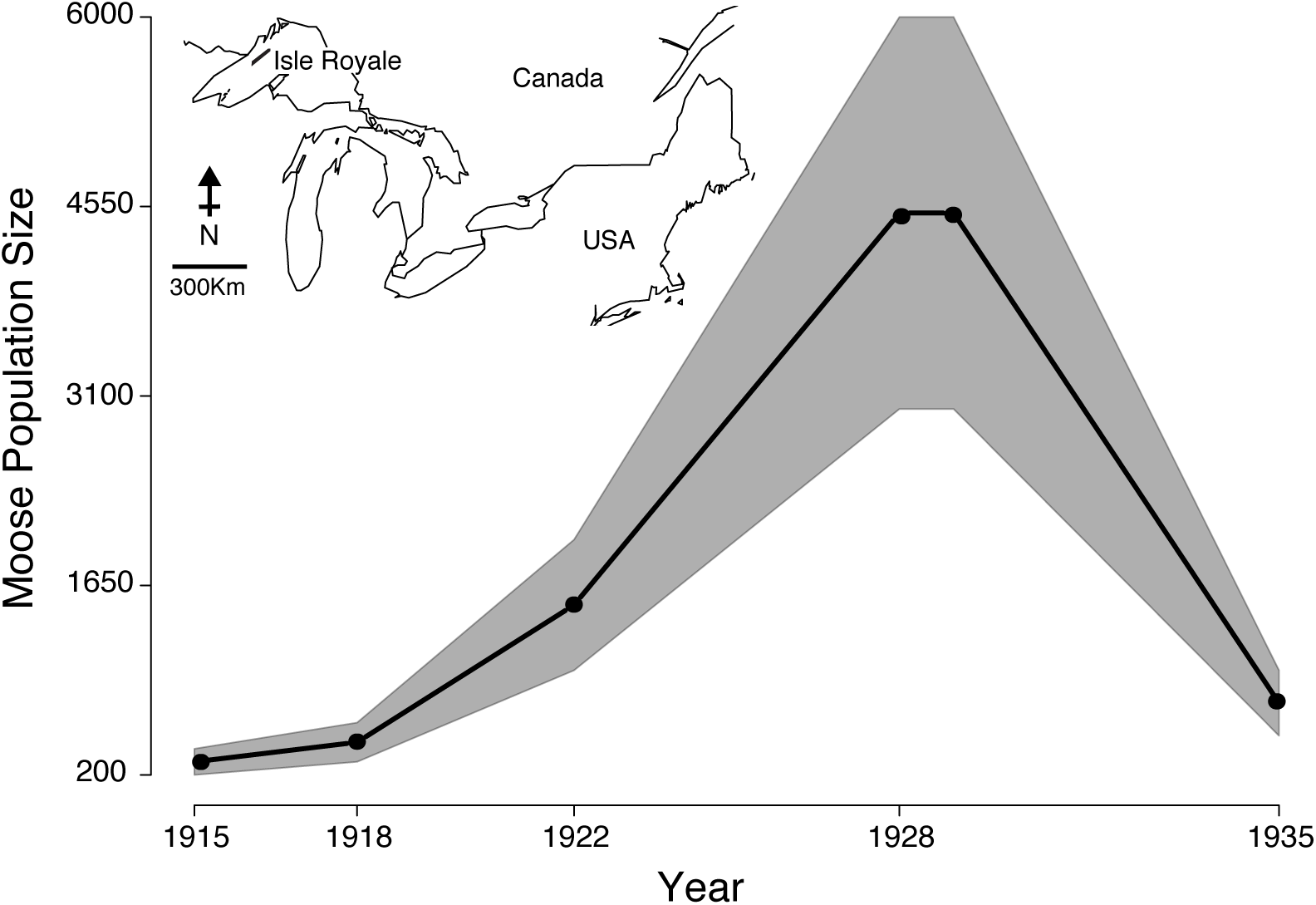
The early population dynamics of moose on Isle Royale. Estimates of moose population size were taken from early naturalist reports of Isle Royale (see map for the island’s location). The points indicate the average trend and the shaded region represents the reported discrepancy. The lower bound of the shaded region represents the actual value reported in the literature and were used for model fitting.

Folklore surrounding the arrival of Isle Royale’s moose speculated that they either swam or crossed over the ice from mainland Canada (Murie, 1934). However, prior to the two-hundred observed in 1915, no moose were recorded on the island and a detailed accounting of the fauna on Isle Royale conducted during 1905 did not report any moose (Adams *et al.*, 1909). Given the large numbers observed in 1915, the relatively narrow time-period in which they arrived, and that moose are solitary animals, e.g. in Minnesota their group sizes rarely exceed seven individuals (Peek *et al.*, 1974), one might question the belief that they swam or walked on to the island (Scarpino, 2011). In fact, the website for Isle Royale National Park now reports that, “Cultural evidence has suggested that a private citizen’s group may have intentionally stocked the moose onto the island for the purposes of recreational hunting (NPS, 2015).” To better evaluate the competing hypotheses surrounding the moose’s arrival on Isle Royale, we fit standard ecological models to data on moose population size from 1915 to 1929. With these models, we estimated the most likely: 1) year of immigration; 2) founder population size; and 3) yearly population growth rate.

Our results suggest that for a logistic growth model, between 40 and 90 moose must have arrived between 1913 and 1914 and subsequently grew at a yearly rate of 0.36–0.43, see Figure 2a. Given that mainland moose near Isle Royale typically live in densities around two moose per square mile (Peek *et al.*, 1971, 1974), this would require all of the moose in between 20 & 45 square miles to band together and travel to Isle Royale. The best-fit exponential growth models required even larger founder population sizes (Figure 2b). Although our estimated growth rate from the logistic model is nearly three times greater than post crash estimates of moose population growth on Isle Royale, see (Messier, 1994), they are in line with estimates of a reindeer population on St. Paul Island, Alaska, USA, which similarly experienced an environment without competitors, predators, human hunting, and with nearly unlimited resources (Scheffer, 1951).

**Figure 2.**
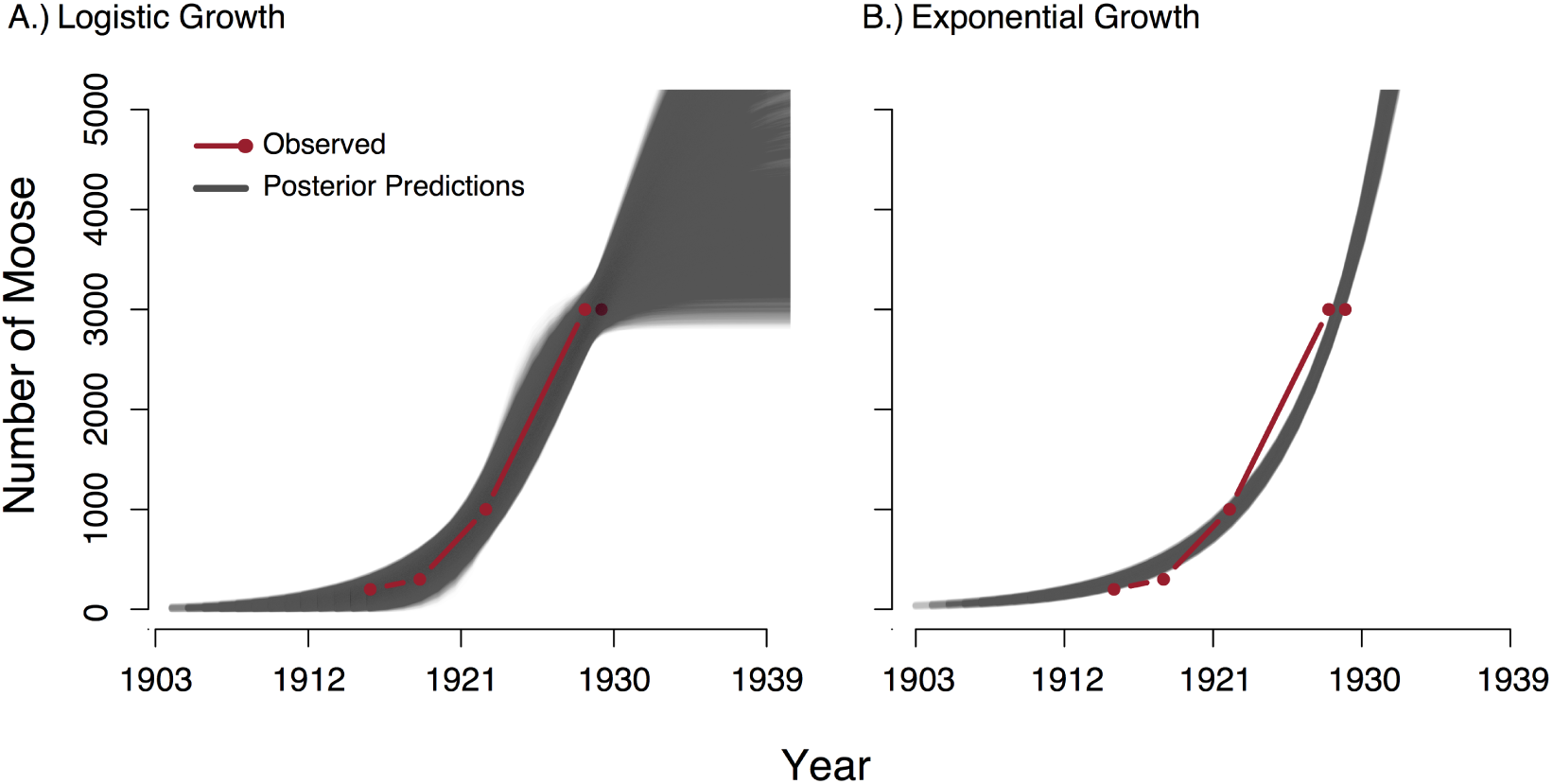
Best-fit growth models for the Isle Royale moose. All parameter sets that were within 6% absolute error were used to plot the moose’s population trajectory (black lines) from: A.) the logistic growth model and b) the exponential growth mode. The observed data used for model fitting are in red.

The population fluctuations observed in the moose of Isle Royale are less dramatic than those seen in the deer population crash on the Kaibab Plateau in Arizona, a crash that was largely the result of United States Forest Service management practices (Binkley *et al.*, 2006). In 1906, the population of deer on the Kaibab Plateau was only a few thousand, by 1921 it had ballooned to over 100,000, and only a few later the deer were headed rapidly towards local extinction. This population increase and crash resulted from the removal of biotic regulation on deer populations; specifically, the interruption of natural fire cycles and the U.S. Forest Service policy of predator reduction (Binkley *et al.*, 2006).

In The Wolves of Isle Royale, Rolf Peterson points out similarly problematic management issues on Isle Royale: “Thus the NPS [National Park Service] policy of maintaining ‘native’ species cannot clearly guide us in our quandary. In an ironic blend of tradition and history, one might argue that neither the wolf nor the moose are purely ‘native’ species at Isle Royale.” Peterson continues his critique of the National Park Service’s non-interventionist policy, stating, “Passive observation, can be an easy policy that doesn’t require much expense or ecological understanding; perhaps that explains some of its appeal. But our national parks deserve better than rote adherence to tradition.”

Our results provide insight into to the early dynamics of the moose population on Isle Royale and represent one of the only estimates of large ungulate population growth in the absence of biotic regulation. Clearly, more population modeling–in combination with population genomic studies of mainland and island moose–would provide greater insight into the origin of Isle Royale’s moose. However, when coupled with the emerging cultural evidence, we must conclude that the moose on Isle Royale may have been introduced by humans. If true, we encourage the National Park Service to re-evaluate its decision not to intervene and save the wolves, and also how they value all of the human cultural resources in our Park system.

## Acknowledgements

The authors acknowledge funding from the Santa Fe Institute and the Omidyar Group to SVS.

## Methods, Data, and Posteriors

### Models and Inference

When populations are self-limiting, their dynamics can be modeled using logistic growth equations. When considering only a single-species, the following differential equation completely describes its population growth over time *t*:

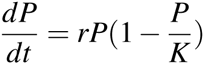

where *P* is the population size, *r* is growth rate, and *K* is the maximum sustainable population size, i.e. the carrying capacity.

The solution for the population size *P* after a certain amount of time *t*, *P*(*t*), with an initial condition that at time *t* = 0 the population had *P*_0_ individuals is:

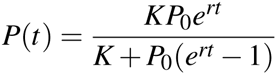

For exponential growth, i.e. growth without a carrying capacity, the corresponding equation for the population size as a function of elapsed time is:

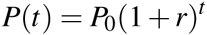

We estimated the parameters of both models from data using a non-linear least squares algorithm coded in the R programming language (R Core Team, 2015). To better explore the uncertainty around these estimated values, we performed Approximate Bayesian Computation using broad, uniform priors over all parameters (Beaumont *et al.*, 2002). Both models were fit to the lower bound on the observed moose population size, clearly a conservative assumption with regard to our conclusions.

### Data

Data were taken from early naturalist reports on Isle Royale as summarized in Paul F. Hickie’s 1934 manuscript, “Isle Royale moose studies,” see Table 1.

**Table 1.** Observed Numbers of moose on Isle Royale 1915 - 1935 as reported in (Hickie, 1936)

| Year | Observed Number of Moose |
| --- | --- |
| 1915 | 200 |
| 1918 | 300 |
| 1922 | 1000 |
| 1928 | 3000–5000 |
| 1929 | 3000–5000 |
| 1935 | 200–500 |

### Posteriors

**Figure 3.**
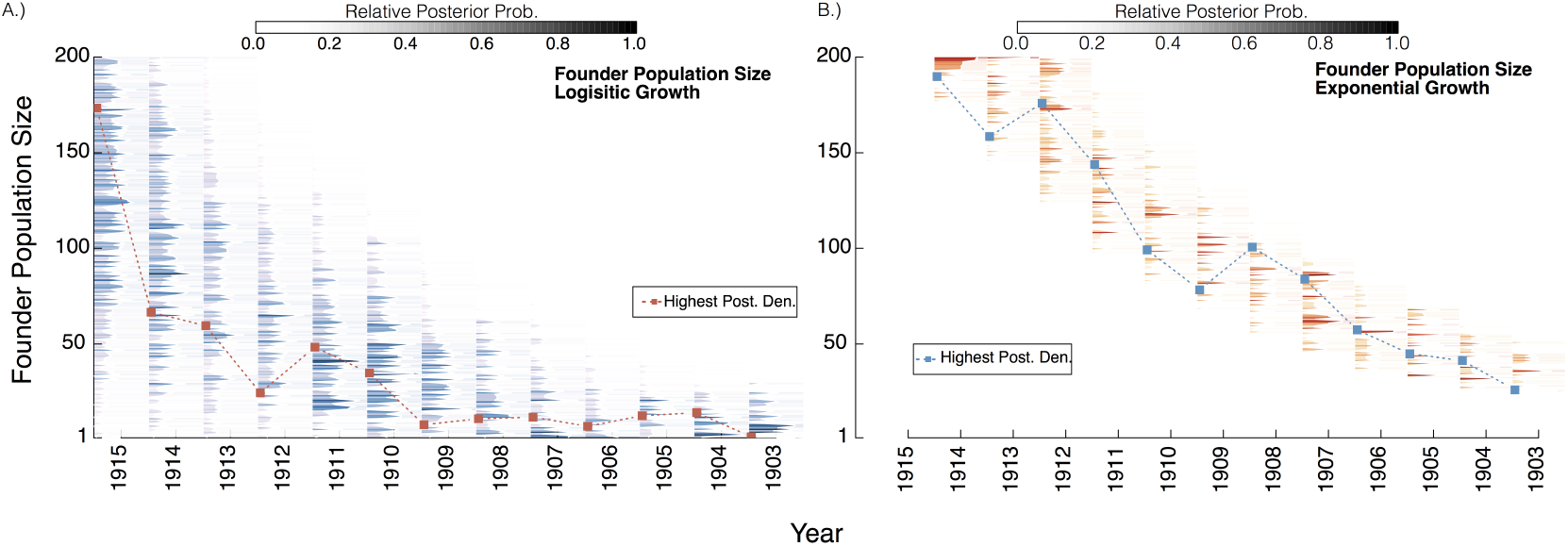
Posterior distribution of the founder population size. The posterior distribution of founder population size is plotted as a function of immigration year for: A.) the logistic growth model and B.) the exponential growth model. All parameters producing fits with 6% absolute error or less were retained, with darker colors indicating progressively better fits.

## References

Adams, C.C., Gleason, H.A., McCreary, O. & Peet, M.M. (1909) An ecological survey of Isle Royale, Lake Superior. Wynkoop Hallenbeck Crawford Company, State Printers.

Beaumont, M.A., Zhang, W. & Balding, D.J. (2002) Approximate bayesian computation in population genetics. Genetics, 162, 2025–2035.

Binkley, D., Moore, M.M., Romme, W.H. & Brown, P.M. (2006) Was aldo leopold right about the kaibab deer herd? Ecosystems, 9, 227–241.

Cronon, W. (1996) The trouble with wilderness: or, getting back to the wrong nature. Environmental History, pp. 7–28.

Cronon, W. (2003) The riddle of the apostle islands. Orion, 22, 36.

Hickie, P.F. (1936) Isle royale moose studies. Transactions of the North American Wildlife Conference, volume 1, pp. 396–399.

Messier, F. (1994) Ungulate population models with predation: a case study with the north american moose. Ecology, 75, 478–488.

Mlot, C. (2015) Inbred wolf population on isle royale collapses. Science, 348, 383–383.

Murie, A. (1934) The moose of isle Royale. University of Michigan Museum of Zoology.

Nelson, M.P. & Callicott, J.B. (2008) The wilderness debate rages on: Continuing the great new wilderness debate. University of Georgia Press.

NPS (2015) Mammals on isle royale-historical context. http://www.nps.gov/isro/learn/nature/upload/Mammals_ver7.pdf. Accessed: 2015-05-05.

Peek, J.M., LeResche, R.E. & Stevens, D.R. (1974) Dynamics of moose aggregations in alaska, minnesota, and montana. Journal of Mammalogy, pp. 126–137.

Peek, J.M., Urich, D.L. & Mackie, R.J. (1971) Moose habitat selection and relationships to forest management in northeastern minnesota. Wildlife Monographs, pp. 3–65.

Peterson, R.O. (1995) The wolves of Isle Royale: a broken balance. University of Michigan Press.

Peterson, R.O. (1999) Wolf-moose interaction on isle royale: the end of natural regulation? Ecological Applications, 9, 10–16.

R Core Team (2015) R: A Language and Environment for Statistical Computing. R Foundation for Statistical Computing, Vienna, Austria.

Scarpino, P.V. (2011) Isle royale national park: Balancing human and natural history in a maritime park. The George Wright Forum, volume 28, pp. 182–198.

Scheffer, V.B. (1951) The rise and fall of a reindeer herd. The Scientific Monthly, 73, 356–362.

